# Validated backbone assignments for natural-abundance ^1^H,^15^N spectra of commercial hen egg white lysozyme and example applications thereof

**DOI:** 10.64898/2026.10.26.761504

**Authors:** Luciano A. Abriata

## Abstract

Although backbone ^1^H and ^15^N assignments for hen egg white lysozyme (HEWL) have been reported in the literature and several related entries are available in the BMRB, these resources differ in sequence, experimental conditions, and the nuclei represented. In particular, there is no BMRB entry with an explicit set of ^1^H,^15^N backbone assignments applicable to spectra that can now be collected at natural ^15^N abundance on concentrated samples of commercial HEWL as shown by Bruker and reproduced here using a 800 MHz magnet with a cryoprobe. Therefore, here I make available the backbone ^1^H,^15^N chemical shifts of commercial (Sigma) HEWL at pH 3.8 and 308 K parsed from previous papers and checked against ^1^H,^1^H NOESY, ^1^H,^1^H TOCSY and ^1^H,^15^N correlation spectra of HSQC and SOFAST-HMQC types, acquired on a 3 mM sample at natural ^15^N abundance. I also provide backbone ^1^H,^15^N assignments transferred to pH 6 and 308 K via a pH titration, closer to physiological conditions as used by others. I discuss details about sample preparation and spectra acquisition, including the measurement times involved, and possible reasons for a few residues lacking assignments. The usefulness of these assignments and of detecting ^1^H,^15^N correlations at natural abundance are proven by (i) application to a titration with a small molecule, (ii) studies of hydration and dynamics via Het-SOFAST-HMQC experiments, and ultimately (iii) by their utilization in published papers even if without attribution. I deliver ready-to-use assignments as text-format Sparky files, and make selected spectra available on the PDB Manipulation Suite’s WebNMR app for straightforward visualization online, useful also for swift educational demonstrations. Overall, I expect the data and methods presented here will be of use in protein biophysics and structural biology research and education, by enabling the interpretation of modern NMR-based studies on inexpensive samples of this widely studied protein.

## Introduction

Hen egg white lysozyme (HEWL hereafter, pictured in Figure 1) is a globular enzyme that has been pivotal in the field of protein biophysics and structural biology due to its simplicity, robustness, and commercial availability at low costs purified from natural sources. Its well-characterized structure, ease of purification, and ready availability in high quality and purity have made it a quintessential model protein, widely used both in advanced research and as a teaching tool in biochemistry and structural biology classrooms. Additionally, HEWL’s ability to hydrolyze bacterial cell walls through cleavage of peptidoglycan makes it an important biological molecule for study, while its thermal stability and compact size (14.3 kDa) have rendered it ideal for various structural techniques, including nuclear magnetic resonance (NMR) spectroscopy.

**Figure 1.**
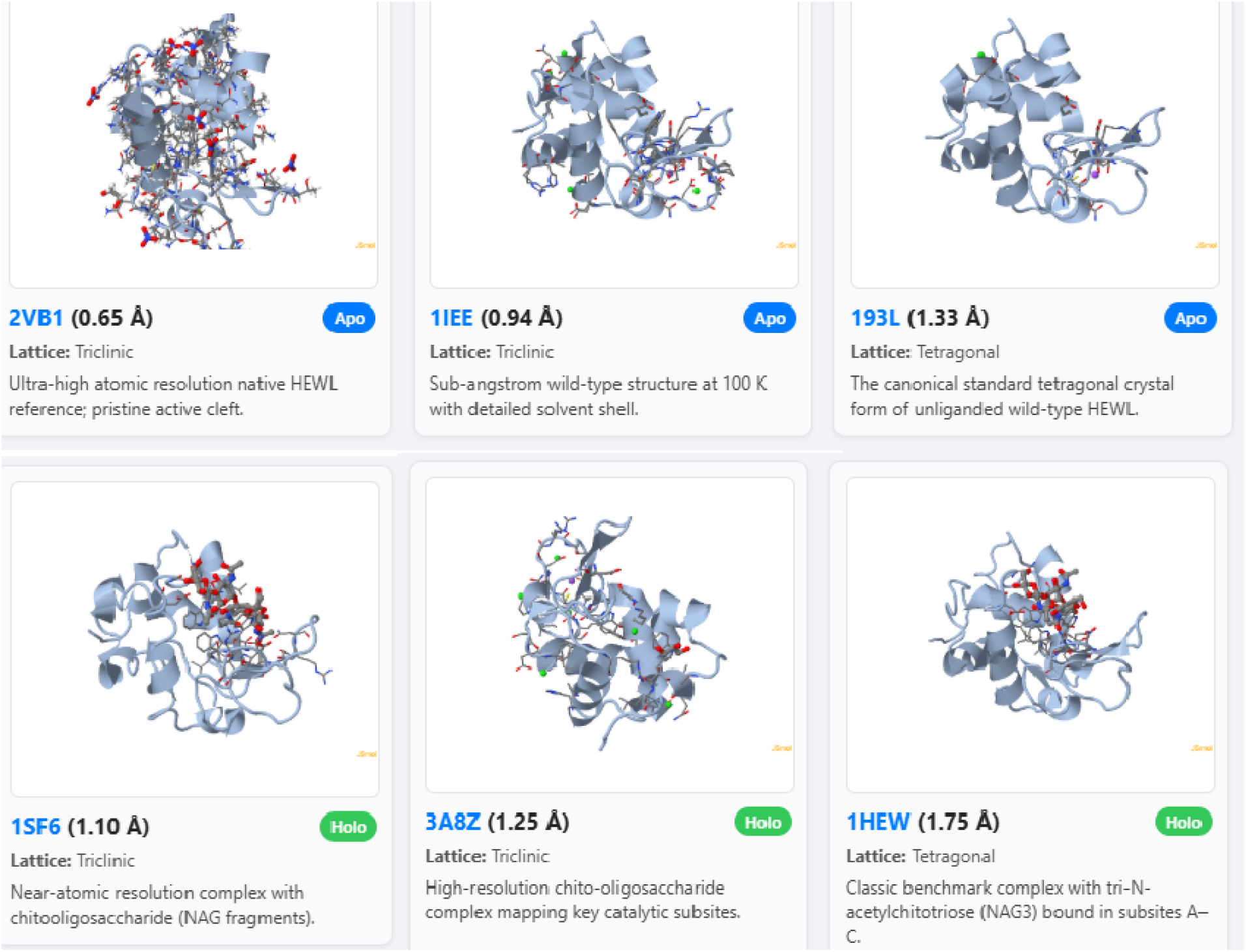
Six higH,resolution PDB entries for hen egg white lysozyme as chosen and rendered automatically by PDBMS-WebMDA’s (https://pdbms.altervista.org/webmda/) copilot,^7^ three in apo form and three loaded with different small molecules. Utilizing the backbone resonance assignments reported here, it becomes possible to study these interactions and many other phenomena via HSQC and SOFAST-HMQC spectra. Interactive 3D versions of the molecular views shown in this figure are available at https://lucianoabriata.altervista.org/lysozyme-natabundance-nmr/hewl-6pdbentries.html

HEWL’s relevance extends to its use as a model system for studying protein folding, dynamics, and ligand binding interactions. Its detailed structural features have been elucidated by X-ray crystallography and NMR, making it an ideal system for studying both its solution-state behavior and solid-state properties. But naturally, it is virtually impossible to obtain HEWL from the natural source in ^15^N-enriched form, and recombinant protein preparation in bacteria requires refolding from urea-denatured protein due to the various intramolecular disulfide bonds that stabilize the protein, a laborious and low-yield protocol.

### Previous NMR assignments of hen egg white lysozyme

HEWL has a long history as a model protein for NMR spectroscopy. Redfield and Dobson reported sequential proton resonance assignments and secondary-structure information for HEWL in 1988.^1^ Buck et al. subsequently reported backbone ^15^N relaxation measurements and associated nitrogen chemical-shift assignments using isotopically enriched lysozyme produced in *Aspergillus niger*.^2^ These studies established a foundation for subsequent investigations of HEWL structure and dynamics.

Several relevant assignment datasets have since become available through the BioMagResBank (BMRB^3^). Entry 1093 contains the historical proton assignments reported by Redfield and Dobson.^1^ Entry 4562 contains extensive ^1^H and ^13^C assignments for native HEWL purchased from Sigma, measured at natural isotopic abundance.^4^ The associated study by Wang et al. reported extensive proton and carbon assignments and discussed the previously published backbone ^15^N assignments. Entry 4943 provides proton chemical shifts for commercial Sigma and Seikagaku lysozyme preparations at pH 3.8 and 308 K.^5^

BMRB entry 4831 contains backbone chemical-shift assignments, including ^15^N shifts, for a recombinant hen lysozyme that contains a Q41E substitution relative to canonical HEWL.^6^ Although many assignments from this entry may be transferable, the sequence difference warrants consideration when interpreting spectra of wild-type commercial protein.

Despite the availability of these resources, the information is distributed across multiple publications and database entries, with differences in sequence, experimental conditions, and the nuclei represented. In particular, the existing resources do not by themselves provide the experimentally evaluated, ready-to-use assignment lists and corresponding natural-abundance ^1^H–^15^N correlation spectra at both pH 3.8 and pH 6.0 that are presented here.

Therefore, in summary, as of October 2026 there is no BMRB entry making the backbone ^1^H,^15^N assignments of commercial HEWL available for use. Some publications have utilized such assignments though, without citing or acknowledging their origin. The BMRB hosts assignments for lysozyme from yet other organisms, but they differ at various residues and hence these assignments cannot be transferred to HEWL spectra in a straightforward and reliable manner. This all represents an unfortunate, quite significant gap in the availability of NMR data for such an extensively studied protein. This more so in light of modern developments in NMR spectroscopy; in particular, of the recent reports showing that it is possible to obtain higH,quality ^1^H,^15^N correlation spectra of commercial HEWL using natural ^15^N abundance in as fast as one hour.^8^

In this work, I compile previously published assignments, evaluate them against two-dimensional NMR spectra acquired on commercial HEWL at natural ^15^N abundance, and transfer the assignments to pH 6.0 through a monitored pH titration. I provide the resulting peak lists in machine-readable and Sparky-compatible formats, document ambiguous and incompletely confirmed assignments, and make representative spectra available for interactive inspection. The aim is to make established HEWL assignment information more directly accessible for routine biophysical research and NMR-based structural biology education.

## Materials and Methods

Lysozyme (>98% purity from hen egg white, without isotopic enrichment, from Sigma, various lots purchased by two other groups at EPFL; see Acknowledgements) was dissolved in TAP buffer (50 mM of each of tris, acetate and phosphate) prepared in 95% H□ O and 5% D□ O, at a concentration of 3-5 mM. The pH was adjusted as required in each case. The sample was subjected to NMR analysis at the indicated pH values and temperatures, using a Bruker 800 MHz spectrometer equipped with a cryogenically cooled probe and an Avance Neo console. For reference, the 90° hard pulse was always around 13-15 µs in this spectrometer (at EPFL’s NMR center, Switzerland) whose hard pulse in pure water with 5% D_2_O and no salts is typically around 8-9 µs.

As indicated in each case, the ^1^H,^15^N correlation spectra were acquired using either the HSQCETF3GP pulse sequence with a D1 of 1s, or SOFAST-HMQC^9^ experiments with a D1 of 0.2 s, in both cases using the pulse sequence available in Bruker Topspin 4 or higher. Het-SOFAST-HMQC spectra^9^ were acquired and setup using NMRlib.^10^ Spectra were acquired and processed using TopSpin 4 or higher, and analyzed using Sparky,^11^ CARA, and the WebNMR app at the PDB Manipulation Suite.^7^ Spectra to transfer and verify assignments (HSQC, NOESY and TOCSY experiments) were carried out at 308 K on samples at pH 3.8. They were then transferred to 308 K and pH 6 via pH titrations monitored by SOFAST-HMQC, with pH points at 3.8, 4.6, 5.2, 5.4, 5.6, 5.8 and 6.0. Covering the range from 5.2 to 6 in increments of just 0.2 pH units was important given some large shifts probably induced by protonation equilibria taking place around pKa 5-6. Of note, the increasing backbone HN exchange at higher pH values was notable, especially so towards the higher pH end (pH 5.8 and 6.0) where longer acquisition times were required (reaching 6-12 horus) and preventing acquisition at pH much higher than 6.5 within reasonable experiment times.

## Results and Discussion

### HSQC and HMQC spectra in the originally utilized conditions

Figure 2 presents the ^1^H,^15^N HSQC spectrum of HEWL as purchased from Sigma at pH 3.8 (in TAP buffer) and 308 K, with all assignments parsed from the two indicated publications that could be verified via 2D TOCSY and 2D NOESY. The well-resolved crosspeaks correspond to backbone amide groups, and the assignments for most residues were confidently transferred from previous studies. The caption to Figure 2 includes a link to download the assignments in Sparky format, and points to a way to interactively explore these spectra online with the WebNMR tool at the PDB Manipulation Suite.

**Figure 2.**
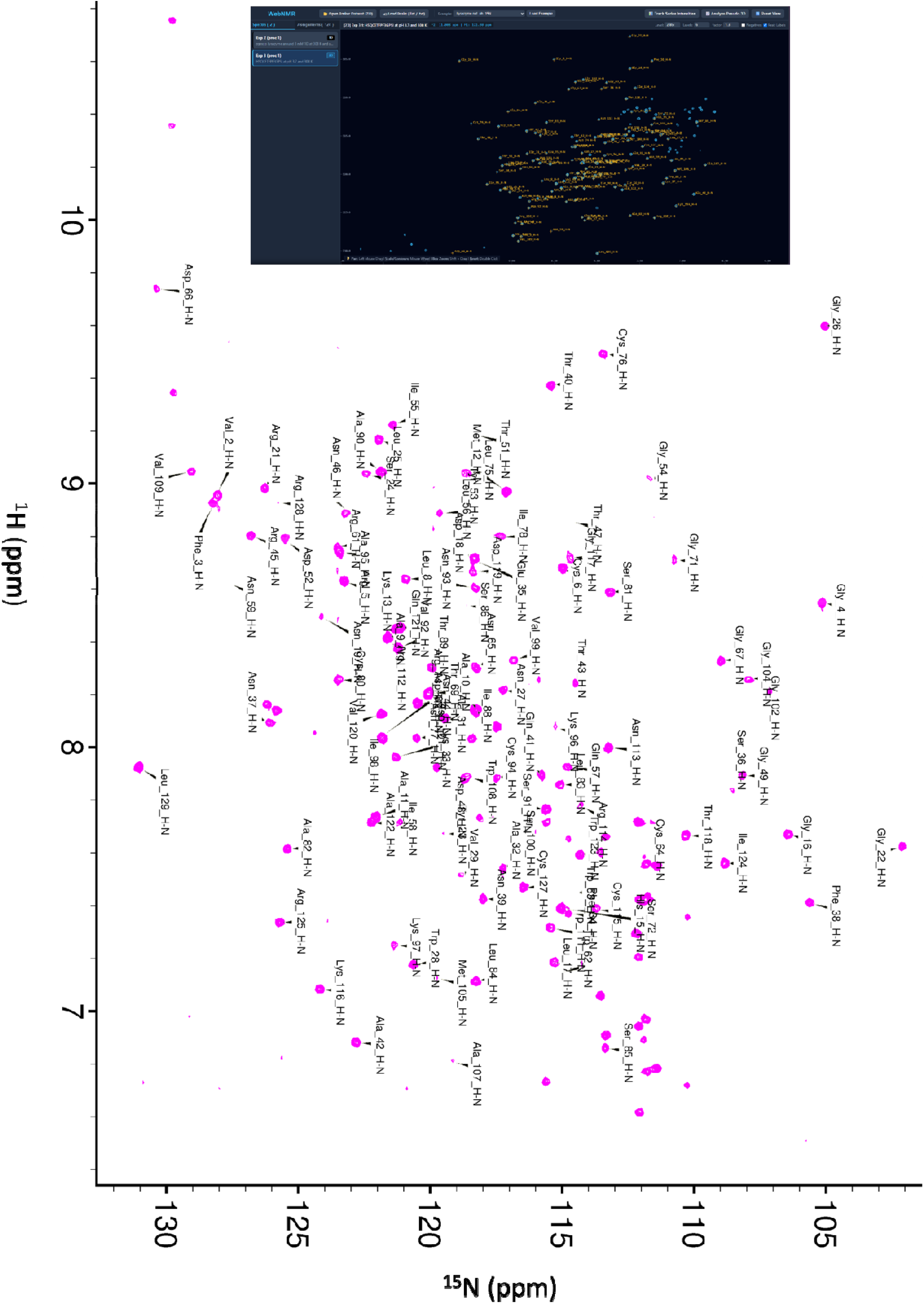
^15^N HSQC of HEWL at pH 3.8, collected in 6 hours at natural ^15^N abundance on a 3 mM sample at 308K, with all assignments parsed from existing papers and verified with 2D TOCSY and NOESY. The assignments are available at http://lucianoabriata.altervista.org/lysozyme-natabundance-nmr/Sigma_HEWL_308K_pH3.8.list. The spectrum can be inspected interactively in experiment number 2 of the HEWL dataset available as example in the WebNMR web app at the PDB Manipulation Suite (screenshot in the inset at the top, https://pdbms.altervista.org/webnmr/).

### pH titration to transfer assignments from pH 3.8 to pH 6, at 308K

In order to achieve assignments at a more physiologically relevant pH, much more useful as demonstrated for example by Mao et al in *Nature* 2025 (who utilized the assignments presented here without explaining their origin nor crediting the original source), I performed a pH titration monitored by 6-12 hour-long SOFAST-HMQC experiments (Figure 3). This resulted in the clean assignment presented in Figure 4 and available for download at the URL indicated in the figure caption.

**Figure 3.**
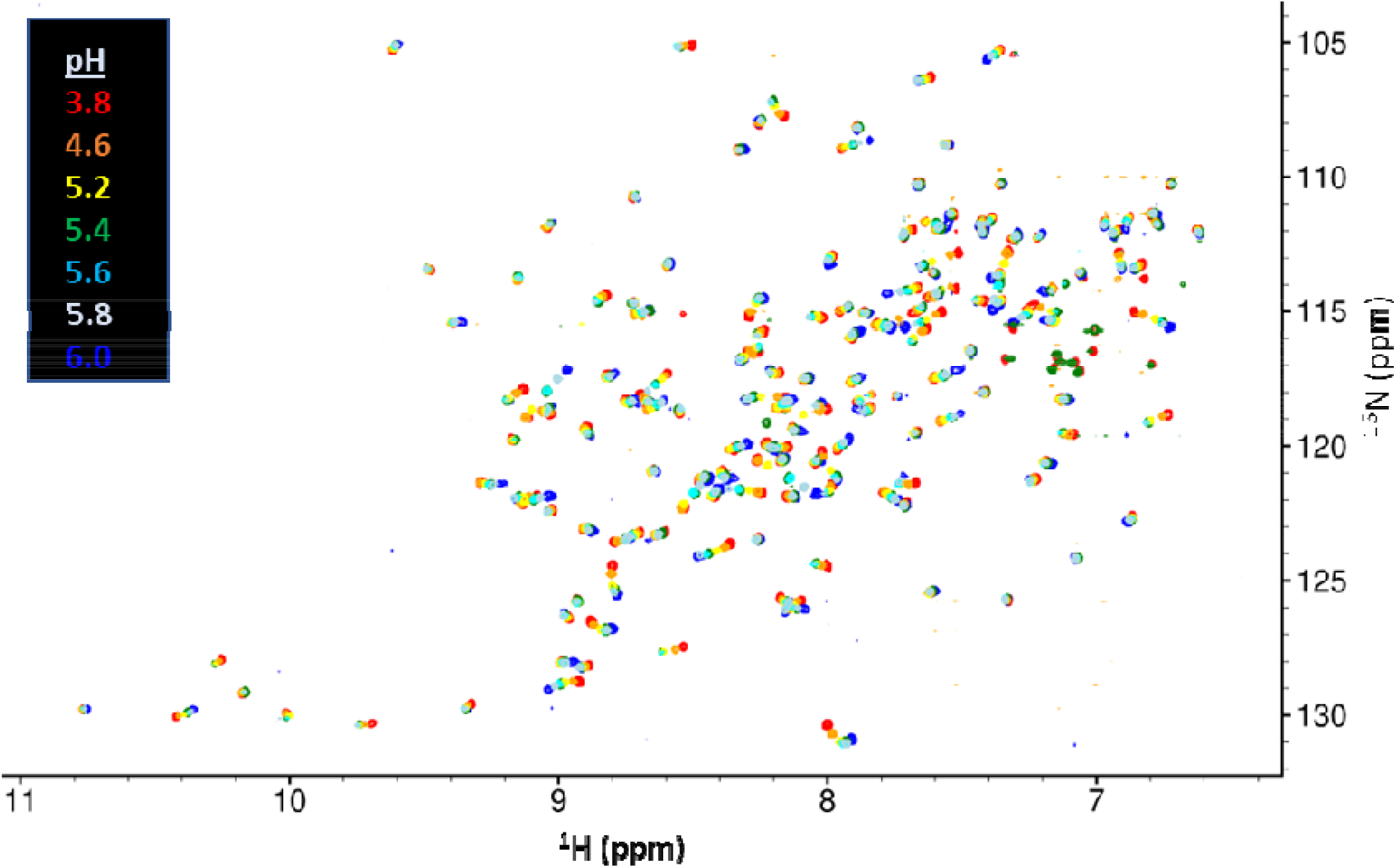
pH titration on HEWL monitored by SOFAST-HMQC spectra at natural ^15^N abundance, at 308 K. pH values are color-coded as in the inset. As a general rule, the higher the pH the longer the required acquisition times, such that in practice the spectra utilized here lasted from 6 to 12 hours. For assignments of the end points, see figures 2 and 4.

**Figure 4.**
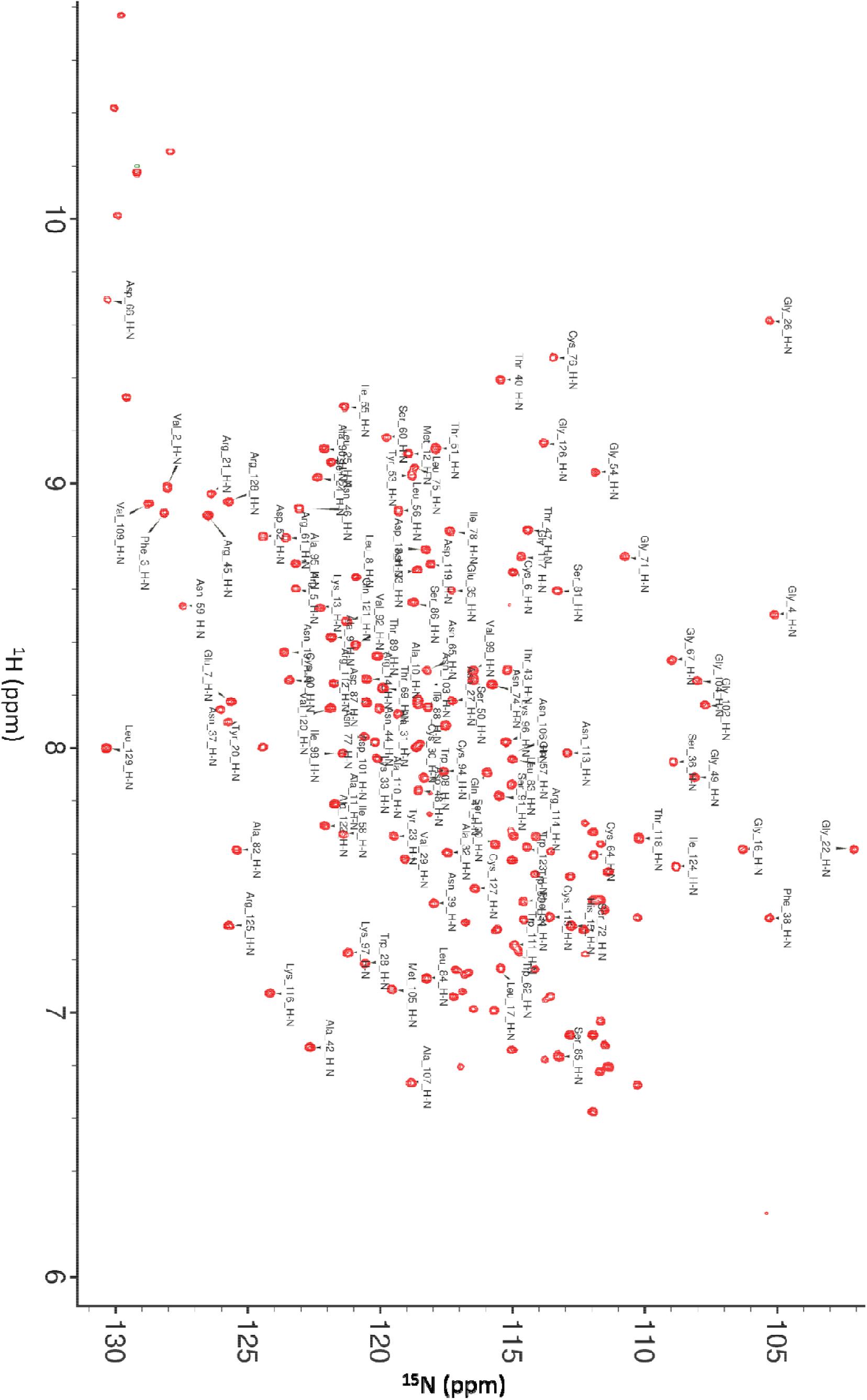
^15^N HSQC of HEWL at pH 6, collected in 12 hours at natural ^15^N abundance on a 3 mM sample at 308K, with all verified assignments transferred from pH 3.8. Assignments are available at http://lucianoabriata.altervista.org/lysozyme-natabundance-nmr/Sigma_HEWL_308K_pH6.0.list

### Notes on the provided assignments

This study provides a curated set of ^1^H,^15^N chemical shift assignments for HEWL as sold by Sigma, at natural ^15^N abundance. This work consolidates and experimentally evaluates previously published HEWL assignments, providing directly usable backbone ^1^H,^15^N peak lists matched to natural-abundance spectra of commercial protein at pH 3.8 and 6.0. By documenting assignment confidence and providing accessible spectral visualizations, the resource complements existing BMRB entries and facilitates the practical use of HEWL in biophysical studies and NMR education.

A note is important regarding the assignments as reported. Ambiguities in certain crosspeaks and exclusions due to insufficient evidence are clearly highlighted, ensuring transparency and usability of the data for downstream research. Especially at pH 6, ambiguity exists for certain residues (C30/A110, L75/Y53, V99/S50, and W62/W111) whose assignments are not included in the table. Additionally, residues N46, Q57, N65, N74, I88, N103, N106, and V120 are only partially confirmed, due to overlap and/or lack of reliable signals and support to back up their assignments. It is notable how many asparagine residues lacked H,N correlations in the spectra, some even at low pH, prompting checks on covalent/primary structure details of the Sigma-supplied HEWL samples via mass spectrometry.

### Example applications

The assignments and methods produced here, particularly the use of SOFAST-HMQC to more rapidly acquire ^1^H,^15^N correlation spectra on concentrated HEWL, were utilized by Mao et al in *Nature* 2025 and are right now serving in at least two more projects at EPFL. To further illustrate their potential applications, Figure 5 shows the results of Het-SOFAST-HMQC^9^ experiments at natural ^15^N abundance (12 hours per spectrum) irradiating the water and aliphatic resonances, with data processed and displayed swiftly at the PDB Manipulation Suite’s WebNMR app. I have made this particular dataset available at WebNMR under the title *Lysozyme nat. ab. Het-SOFAST-HMQC*, here visualized on PDB ID 2VB1. As the reader can verify on the raw spectra interactively at WebNMR, the signal-to-noise ratio is very good; this is reflected in the clear profiles after the data is mapped on the structure, especially for water irradiation where the more exposed and flexible regions display lower values (light blue in Figure 5 center).

**Figure 5.**
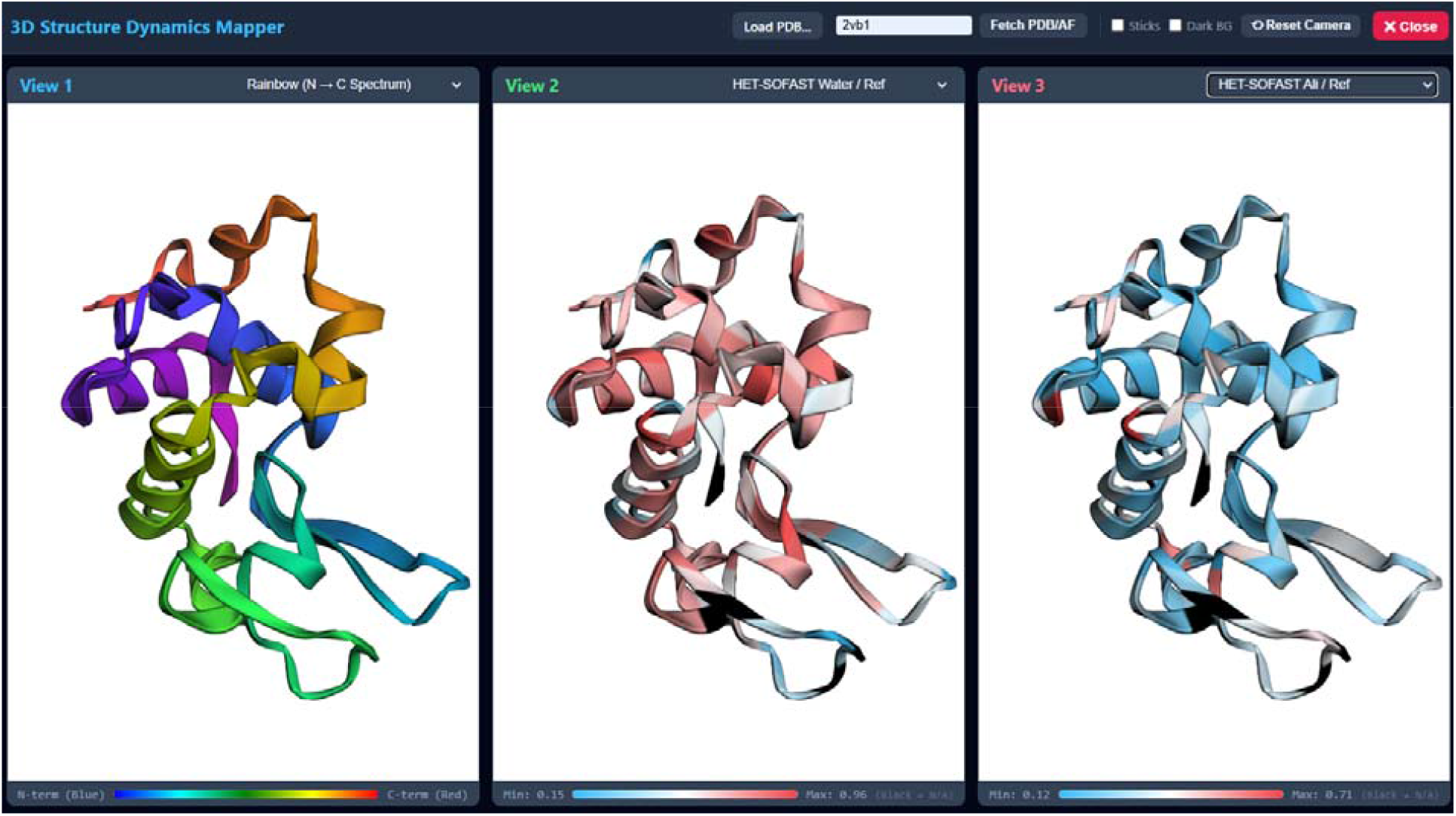
Mapping of Het-SOFAST-HMQC data collected at natural ^15^N abundance on HEWL at pH 6 on a concentrated HEWL sample, at 308 K. These data are available as an example (Lysozyme nat. ab. Het-SOFAST-HMQC) at the WebNMR app of the PDB manipulation suite (https://pdbms.altervista.org/webnmr/), where it was processed and displayed using PDB ID 2VB1.

In turn, Figure 6 shows the results of mapping the binding of a ligand that target a wide area of the protein’s surface, as revealed by titrations monitored by SOFAST-HMQC (analyzed with the WebNMR app at the PDB Manipulation Suite).

**Figure 6.**
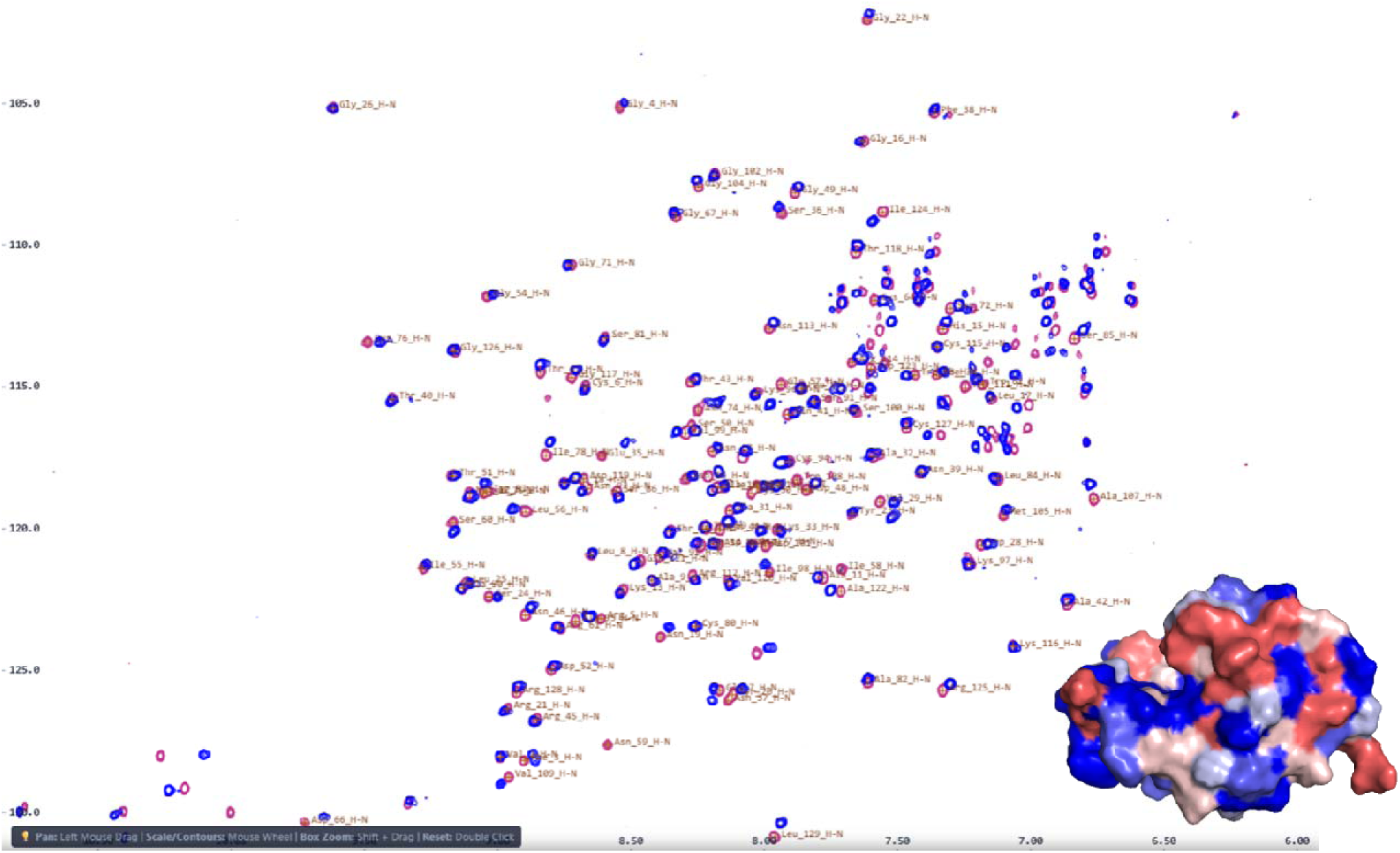
Titration of a small molecule into commercial HEWL, followed by ^15^N SOFAST-HMQC at natural abundance. Free HEWL (pink) and at the end point of a titration with a small molecule that targets a wide surface patch (blue). Spectra were overlaid in the PDBMS’ WebNMR app, applying the assignment at pH 5.2. At the bottom right I show WebNMR’s own mapping of the residues experiencing high minimal CSP, spanning from blue (< 0.02 ppm in blue to > 0.05 ppm in red). The structure is PDB ID 2VB1.

## Conclusion

In summary, this short work provides validated, publicly accessible backbone ^1^H,^15^N assignment tables and Sparky-ready peak lists for commercial HEWL at both acidic (pH 3.8) and near-physiological (pH 6.0) conditions. By demonstrating that higH,field spectrometers equipped with cryogenic probes can deliver higH,quality 2D correlation spectra at natural ^15^N abundance within hours, this study removes the barrier of expensive isotopic labeling for HEWL studies of effects from perturbations on pH, solvent, ligand binding, dynamics, etc. as exemplified with Het-SOFAST-HMQC and titration experiments. These resources will facilitate routine biophysical characterizations, ligand binding screens, and hands-on structural biology laboratory training using an affordable, readily available benchmark protein.

## Data availability

The assignments are available in Sparky format in two separate files for pH 3.8 and 6, at the following Zenodo link: https://zenodo.org/records/23453594. Direct links are http://lucianoabriata.altervista.org/lysozyme-natabundance-nmr/Sigma_HEWL_308K_pH3.8.list and http://lucianoabriata.altervista.org/lysozyme-natabundance-nmr/Sigma_HEWL_308K_pH6.0.list. Some of the spectra presented here, including the Het-SOFAST-HMQC spectra irradiating water and aliphatics at pH close to 6 and comparable to the corresponding no-irradiation reference to reproduce Figure 5, are available as off-the-shelf examples at the PDB-MS’s WebNMR app at https://pdbms.altervista.org/webnmr/

## Acknowledgements

This work was motivated by three separate projects that came in at EPFL’s Protein Structure Core Facility from three labs at around the same time. I therefore thank the people involved (A. Agnook, C. Siri, and J.-R. de Gea) who provided most of the samples utilized here and helped run some of these spectra. I also thank A. Bornet (ISIC, EPFL) for free NMR time at the 800 MHz instrument, without which this work would not have been possible.

